# CAMML with the Integration of Marker Proteins (ChIMP)

**DOI:** 10.1101/2022.05.10.491331

**Authors:** Courtney Schiebout, H. Robert Frost

## Abstract

**Motivation:** Cell typing is a critical task in the analysis of single cell data, particularly when studying diseased tissues that contain a complex mixture of normal tissue and infiltrating immune cells. Unfortunately, the sparsity and noise of single cell data make accurate cell typing at the level of individual cells extremely difficult. To address these challenges, we previously developed the CAMML method for multi-label cell typing of single cell RNA-sequencing (scRNA-seq) data. CAMML uses weighted gene sets to score each profiled cell for multiple potential cell types. While CAMML outperforms other scRNA-seq cell typing techniques, it only leverages transcriptomic data so cannot take advantage of newer multi-omic single cell assays that jointly profile gene expression and protein abundance (e.g., joint scRNA-seq/CITE-seq).

**Result:** We developed the ChIMP (CAMML with the Integration of Marker Proteins) method to support multi-label cell typing of individual cells jointly profiled via scRNA-seq and CITE-seq. ChIMP combines cell type scores computed on scRNA-seq data via the CAMML approach with discretized CITE-seq measurements for cell type marker proteins. The multi-omic cell type scores generated by ChIMP allow researchers to more precisely and conservatively cell type joint scRNA-seq/CITE-seq data.

## Introduction

The immune cells present in tissues and organs have important implications for tissue health and function, particularly in diseased states.^1–3^ Tumor-infiltrating immune cells are an important example, with immune cell presence and phenotype being key indicators of disease outlook and prognosis.^2–7^ In particular, the polarization state of macrophages and the presence and phenotype of T cells can predict the severity of tumor immune evasion and its downstream effects on tumor growth.^7–11^ Thus, identifying infiltrating immune cells and characterizing their cell (sub)type and phenotype is a critical aspect of cancer research, with particular relevance to the analysis of tumor genomic data measured at the bulk or single cell level. For the analysis of bulk tissue data, e.g., bulk RNA-sequencing (RNA-seq), a common approach involves the application of deconvolution methods (e.g. CIBERSORT^12^, DeconRNAseq^13^) to estimate the proportions of each cell type in the tissue. However, bulk tissue analysis only provides estimates of cell type proportions, requires prior knowledge of the cell types present in the tissue, and does not work well for small cell populations or genes with low expression.^14,15^ These limitations inhibit bulk devolution methods from accurately detecting all cell types in a tissue without bias toward the phenotypes that are most highly expressed.

In order to reduce this lack of granularity and specificity, utilization of single cell RNA-seq (scRNA-seq) has become increasingly popular for characterizing tissues.^2,7,8,16,17^ This allows each cell’s individual transcriptome to be analyzed independently of the other cells present in the tissue, allowing for smaller signals and cell populations to be detected. However, this approach is not without limitations with noise and sparsity key challenges for the analysis of scRNA-seq data.^18,19^ To overcome these issues, cell typing of scRNA-seq data is usually performed at the cluster level rather than for single cells.^20–22^ Specifically, it is common practice for investigators to cluster scRNA-seq data, determine the most differentially expressed (DE) genes present in each cluster, and then manually deduce the most likely cell type identity of each cluster based on the DE genes.^21^ While this method leverages known biological context to identify cell types, it makes the assumption that all cells within a cluster have the same cell type, which is often not the case. Furthermore, given that cells are clustered by an unsupervised algorithm that considers all expressed genes, cells that are phenotypically similar may be assigned the same cell type even when the underlying cell identities are distinct (e.g., cytotoxic NK and T cells).^21,23^ Cell-typing at the cluster level can therefore lead to a high rate of misclassification, especially among phenotypically similar cell types.

Methods that assign cell types to individual cells, and thus do not assume cluster homogeneity, have been developed in response to this issue, often using correlation or cell similarity to identify individual cell types.^24,25^ However, these methods were designed to identify just a single cell type for each cell, despite growing evidence that many cell types, especially immune cell types, occur on a continuum, rather than in discrete categories.^9,11^ To address this limitation, we developed cell typing using variance Adjusted Mahalanobis distances with Multi-Labeling (CAMML), a multi-label scRNA-seq cell typing method that utilizes weighted cell type gene sets to score cells for their most likely identities.^26^ We found that CAMML achieved classification performance that was equal to or superior than existing methods with the added benefit of characterizing cells whose phenotype is on a spectrum between standard cell types.^26^ These features allow CAMML to better capture the underlying biology of complex cell populations where there is phenotypic overlap between cell types and differentiation-driven changes in cell state. Although CAMML provides a number of advantages relative to existing scRNA-seq cell typing techniques, it only utilizes transcriptomics data for cell type estimation so is unable to fully leverage the information generated by new multi-omic single cell assays.

Advances in single cell profiling techniques now allow investigators to measure multiple omics modalities on each cell. One such modality being employed in combination with scRNA-seq is Cellular Indexing of Transcriptomes and Epitopes by sequencing (CITE-seq), which quantifies the abundance of cell surface markers in single cell analysis.^27^ This allows cell type-relevant protein markers on the surface of cells, such as CD4 and CD8 in T cells, to be identified and quantified at a single cell level.^27^ Given that cells are often identified by their surface markers, this technology is inherently informative for cell typing of single cell data. To leverage both surface protein and transcriptomic information for multi-label cell typing, we developed CAMML with the Integration of Marker Proteins (ChIMP), an extension of CAMML that allows users to perform cell typing of joint CITE-seq/scRNA-seq data. ChIMP follows the basic framework outlined in CAMML for cell type scoring with scRNA-seq data with an integration step for joint CITE-seq measurements to provide a conservative, highly specific filter for cell classification.

## Methods

### 1. Single cell data sources and processing pipeline

Data was accessed from a study by Lawlor, et al.^28^ where peripheral blood mononuclear cells (PBMCs) was isolated from healthy individuals and split into 2 samples.^28^ One sample was used to perform flow cytometry and the other sample was analyzed with joint scRNA-seq and CITE-seq. Any cells labeled by the original paper as multiplets were removed prior to further analysis.^28^ Only cells with at least 1,000 genes, of which no more than 5% were mitochondrial, were kept for analysis. Furthermore, any genes present in fewer than 100 cells were removed. The CITE-seq was merged following the filtering of the scRNA-seq data but no specific CITE-seq filtering steps were applied. The scRNA-seq was then log-normalized and the CITE-seq was normalized by center log ratio transformation and both omics modalities were scaled.^23^ Following these filtering steps, 37,048 cells with 13,764 genes were left for downstream analysis. These cells were then visualized using UMAP^29^ on 30 dimensions and clustered with a resolution of 0.25 using the Louvain algorithm^30^ as implemented in the Seurat framework, resulting in 9 clusters.^23^ An additional joint scRNA-seq and CITE-seq dataset of PBMCs with 228 surface protein antibodies from HIV vaccine trial patients in Hao, et al.^31^ was filtered, scaled, and normalized in the same way as the aforementioned dataset. Following this processing, 56,775 cells with count data for 17,808 genes remained, separating into 19 clusters. The final dataset utilized for evaluation was joint scRNA-seq/CITE-seq of Mucosa-Associated Lymphoid Tissue (MALT) B cell tumors.^32^ Because this dataset only profiled about 9500 cells, a less stringent Seurat filtering pipeline was applied to maintain cell numbers.^23^ Specifically, only cells with at least 100 genes with nonzero counts, of which no more than 10% were mitochondrial, were kept for analysis, and any genes present in fewer than 10 cells were removed. These cells were then scaled and normalized in the same way as the other two datasets, but were visualized using UMAP^29^ on only 10 dimensions and clustered with the Louvain algorithm^30^ in Seurat with a resolution of .5.^23^ Following this, the dataset contained 6,438 cells in 11 clusters.

### 2. CAMML method

To ameliorate the notable challenges of noise, sparsity, and continuous phenotypes for cell typing scRNA-seq data, we previously developed cell typing using variance Adjusted Mahalanobis distances with Multi-Labeling (CAMML).^26^ CAMML leverages cell type gene sets through Variance-Adjusted Mahalanobis (VAM)^33^ distance scoring to provide single or multi-label cell typing of scRNA-seq data. The Variance-Adjusted Mahalanobis (VAM) method computes cell-level gene set scores using the squared value of a modified Mahalanobis multivariate distance that is measured from the origin rather than the multivariate mean and adjusts for only the technical variance of the genes in each set rather than the full sample covariance matrix.^33^ These squared modified Mahalanobis distances are computed on both the original scRNA-seq data and data where the cell labels are permuted to break inter-gene correlation. To determine a null distribution for these distances, a gamma distribution is fit to the non-zero distances for the permuted data. Scores for each cell are computed using this null distribution by finding the cumulative distribution function (CDF) value for the non-permuted squared distances resulting in a score from 0 to 1 for each cell. These scores are robust to large values, can be compared between gene sets of different sizes and can be subtracted from 1 to generate valid p-values under the null of uncorrelated technical noise.^33^ To allow for prioritization of particular genes in a given set, we updated VAM to accept positive gene weights.^26,33^ These gene weights are used to modify the technical variance for each gene by dividing the original technical variance estimate by the associated gene weight.^26^ This adjustment prioritizes genes with large weights by reducing the associated variance penalty in the modified Mahalanobis distance computation. For the CAMML method, gene-level weights are used to prioritize genes that are highly characteristic of a cell type (i.e. CD8a in CD8+ T-cells) in the computation of VAM scores for cell type gene sets.^26,33^ Support for gene weights is included in the 1.0.0 version of the VAM R package, which is available on CRAN.

Building on the weighted version of VAM, we developed the CAMML method and associated R package for multi-label cell typing of scRNA-seq data.^26^ CAMML uses the weighted version of VAM to compute cell-level scores for gene sets representing cell types. These cell type-specific gene sets are defined to include both genes differentially expressed (DE) in the cell type as well as genes annotated to one of the MSigDB C8^34^ gene sets associated with that cell type. The inclusion criteria and associated weight for DE genes is based on either the log-fold change in expression in the target cell type relative to other cell types or the -log(p-value) from the edgeR^35^ DE test. These genes are then intersected with the associated MSigDB C8 cell-type collection to designate the final gene set. These cell type gene sets can then be scored for a query scRNA-seq dataset using the weighted VAM algorithm outlined above and these scores can be leveraged to classify cells as desired.^26,33^

### 3. ChIMP method

To support cell typing of joint scRNA-seq/CITE-seq data, we have developed a simple but robust method for integrating CITE-seq data into the original CAMML method. We have named this technique “CAMML with the Integration of Marker Proteins (ChIMP)”. As visualized in Figure 1, the ChIMP method performs multi-label cell typing of joint scRNA-seq/CITE-seq data using the following steps:

**Figure 1.**
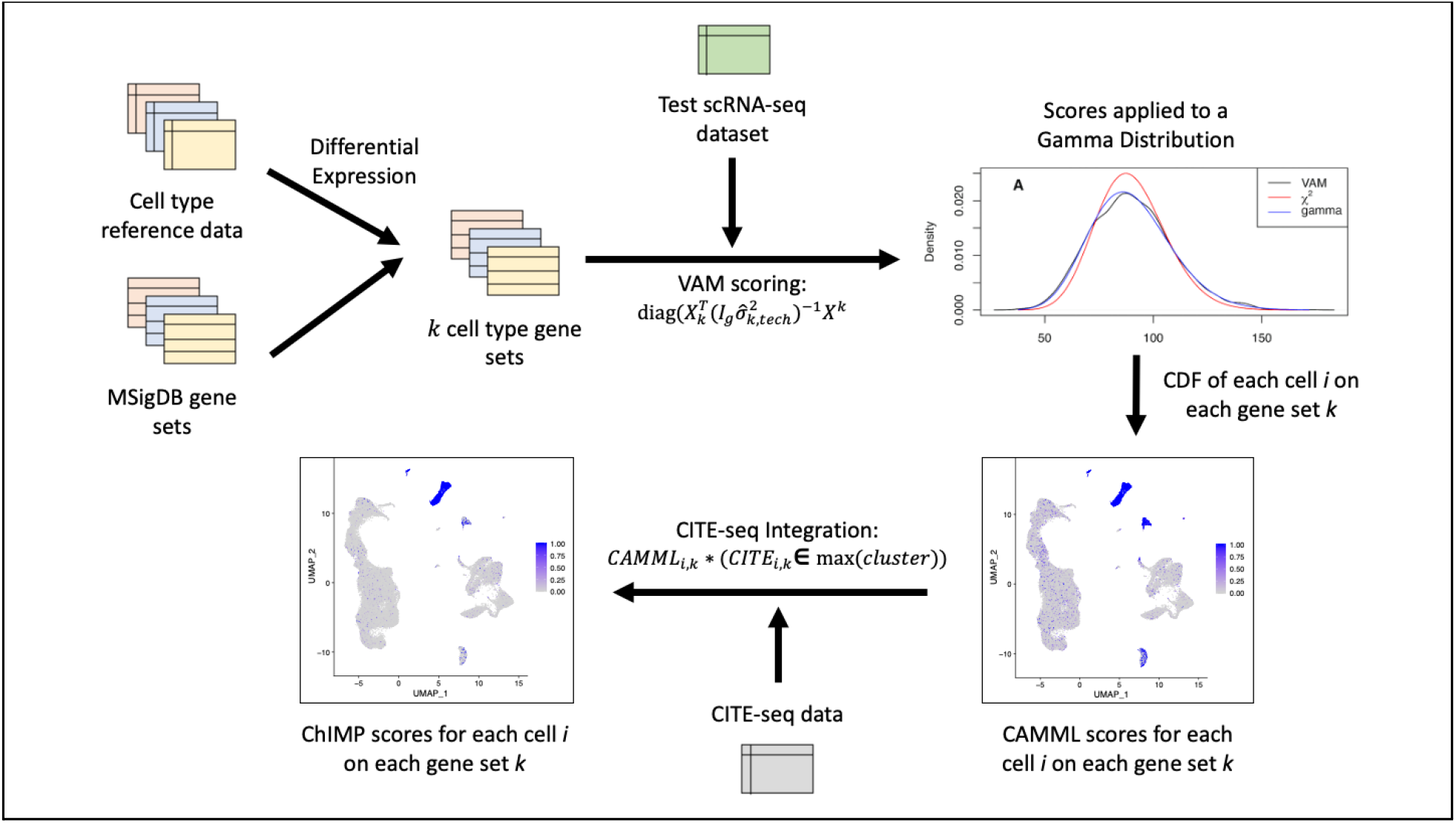
ChIMP pipeline. scRNA-seq data undergoes VAM scoring based on reference cell type gene sets to receive CDF scores for cell types. These scores are then modified by the integration of CITE-seq data to produce ChIMP scores, visualized for B-cells on the Hao, et al. data.^31^

1. For each cell type surface protein marker profiled via CITE-seq, k-means clustering of the CITE-seq count data with 2 centers is used as a method for binarization. In other words, if a marker’s count number in a given cell is in the lower count values cluster, the CITE-seq score becomes 0; if it is in the higher count values cluster, the CITE-seq score will be 1. In cases where more than one marker is sufficient for a cell type (i.e. CD4 and CD8 in T cells), if either marker is in the high value cluster, the score is assigned a 1. If neither marker is in the high value cluster, the score is 0. The discretization of k-means clustering was selected for its ability to robustly discern between the typically bimodal distributions of CITE-seq counts. When compared to another discretization option in the form of the median, the cut-off between k-means clusters proved to be more effective at discerning between the two peaks (Figure 2).

**Figure 2.**
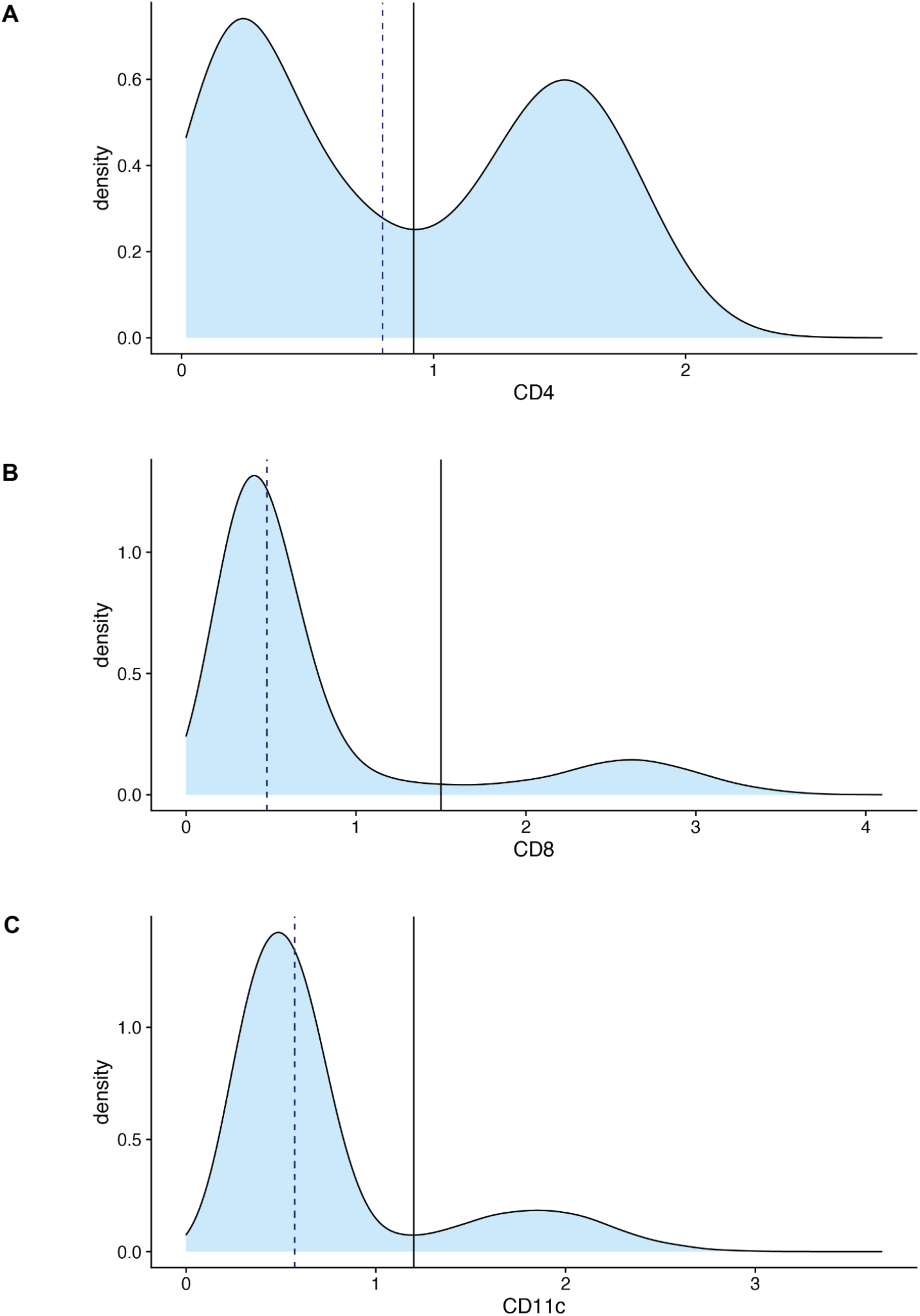
CITE-seq Distributions. A visual of the densities of CITE-seq counts for three example markers **A**. CD4, **B**. CD8, and **C**. CD11c with the median marked with a dashed line and the k-means clustering cut-off marked with a solid line.
2. The discretized cell type CITE-seq scores for each cell are then multiplied by the associated CAMML scores, resulting in an overall cell type score of 0 if the CITE-seq count is in the lower count values cluster for a given marker, and maintaining the original CAMML score if the CITE-seq count is in the higher values cluster.

It is important to note that the technique used by ChIMP to integrate CITE-seq data can never increase the scores computed using just scRNA-seq data. This makes ChIMP a strictly conservative modification of the original CAMML method, i.e., it will only lower sensitivity and increase specificity for cell type classification. If the generated scores are transformed into p-values and used for inference, the ChIMP method will result in a more conservative test, i.e., the type I error rate will be equal to or lower than the nominal alpha.

### 4. Comparative cell typing methods

We performed a comparative evaluation of ChIMP against the original CAMML method, cell typing based on continuous CITE-seq values, cell typing based on discretized CITE-seq values, and SingleR.^25^ CAMML cell typing was done using gene sets built from genes with a logFC greater than 5 in edgeR^35^ DE analysis of the Human Primary Cell Atlas (HPCA).^25,36^ The DE genes were then intersected with the bone marrow cell type gene sets available in the C8 collection of the Molecular Signatures Database (MSigDB), version 7.5.1.^17,34^ These cell type gene sets were then scored with VAM, weighted by the logFC of each gene, resulting in cell-level scores for each supported cell type.^26,33^

Continuous CITE-seq cell typing was performed to give each cell a score from 0 to 1 based on the abundance of surface protein markers for each cell type. This was accomplished by creating an empirical CDF (eCDF) for all the normalized and scaled CITE-seq counts for each cell surface marker.^37^ Individual cells were then scored based on the eCDF value associated with the cell-level marker abundance. This approach generates scores on the same scale as those generated by CAMML and ChIMP. In cases where a single cell type label was needed, the cell type whose surface marker had the highest eCDF score was used. Throughout this manuscript, this method will be referred to as CITE-seq eCDF.

Discretized CITE-seq cell typing was also used as a comparative approach given that discretization is utilized for the integration of CITE-seq data in ChIMP. For this approach, each cell is given a binary score for each cell type based on whether the CITE-seq count for a given surface marker belongs to the lower or higher value cluster of CITE-seq counts for that surface marker across all profiled cells.

Lastly, SingleR^25^ was used as an independent comparative measure for cell typing accuracy. HPCA^36^ was used as the reference for this method using the following cell types: B cells, NK cells, T cells, and monocytes. The single cell type labels called by SingleR were used for accuracy and cell proportion comparisons.^25^

### 5. Entropy analysis

To evaluate ChIMP cell typing specificity, we leveraged a method for measuring entropy previously applied in the original CAMML manuscript: modified Shannon Diversity Index (mSDI), which is defined in Equation 1.^26^ This statistic allows cells to be scored based on both the strength of their ChIMP scores and the number of how many cells are present. In ordinary SDI, if two cells only had nonzero ChIMP scores for T cells (i.e., the scores for all other evaluated cell types are 0), but those T cell scores differed, they would still have identical entropy values. However, by modifying the SDI, the cell with a higher T cell score receives a lower entropy score.

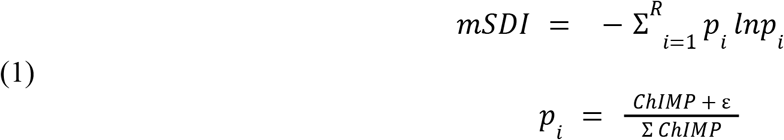

### 6. Differential expression

To better understand cases of discordant cell typing across transcriptomic expression and marker protein abundance, VAM analysis was performed on the Hao dataset for all Kyoto Encyclopedia of Genes and Genomes (KEGG)^38^ gene sets available in version 7.5.1 of MSigDB.^34^ VAM CDF scores for each KEGG pathway were then compared by differential expression analysis between clusters via Wilcoxon Rank Sum test within Seurat’s “FindAllMarkers” function.^23,39^ The log fold change (logFC) threshold between groups was required to be greater than .01 and only upregulated VAM pathways were considered. The same functions were also run on both the gene expression data and CITE-seq surface marker expression data with identical parameters.

## Results and Discussion

### 1. Lawlor Flow Cytometry data

To compare ChIMP performance relative to CAMML, CITE-seq eCDF, and SingleR, we analyzed the joint scRNA-seq/CITE-seq and flow cytometry dataset compiled by Lawlor, et al.^28^ To generate this dataset, flow cytometry and joint scRNA-seq/CITE-seq were performed in parallel on each experimental sample. While this approach does not provide ground truth cell type labels for the single cell data, it allows the cell type proportions computed on the single cell data to be compared to the proportions measured by flow cytometry. The original paper performed FAC sorting to manually discern 4 cell types: B cells, monocytes, NK cells, and T cells. For the scRNA-seq/CITE-seq data, SingleR^25^, CAMML and ChIMP were used to estimate cell types. For ChIMP, the transcriptomic method from CAMML was integrated with CITE-seq markers for each cell type: CD19 for B cells, CD14 for monocytes, CD56 for NK cells, and either CD4 or CD8 for T cells. The proportions of each cell type from SingleR, single-label CAMML, single-label CITE-seq eCDF, and single-label ChIMP were then compared for accuracy. In every case, the cell label proportions were highly correlated with the flow cytometry proportions, ranging from .88 to .95 (Figure 3). The top cell type label based on the eCDF of each CITE-seq marker was the most correlated with the flow cytometry proportions, which is not surprising given that CITE-seq values are based on abundance of the same receptor-specific antibodies used for FAC sorting.

**Figure 3.**
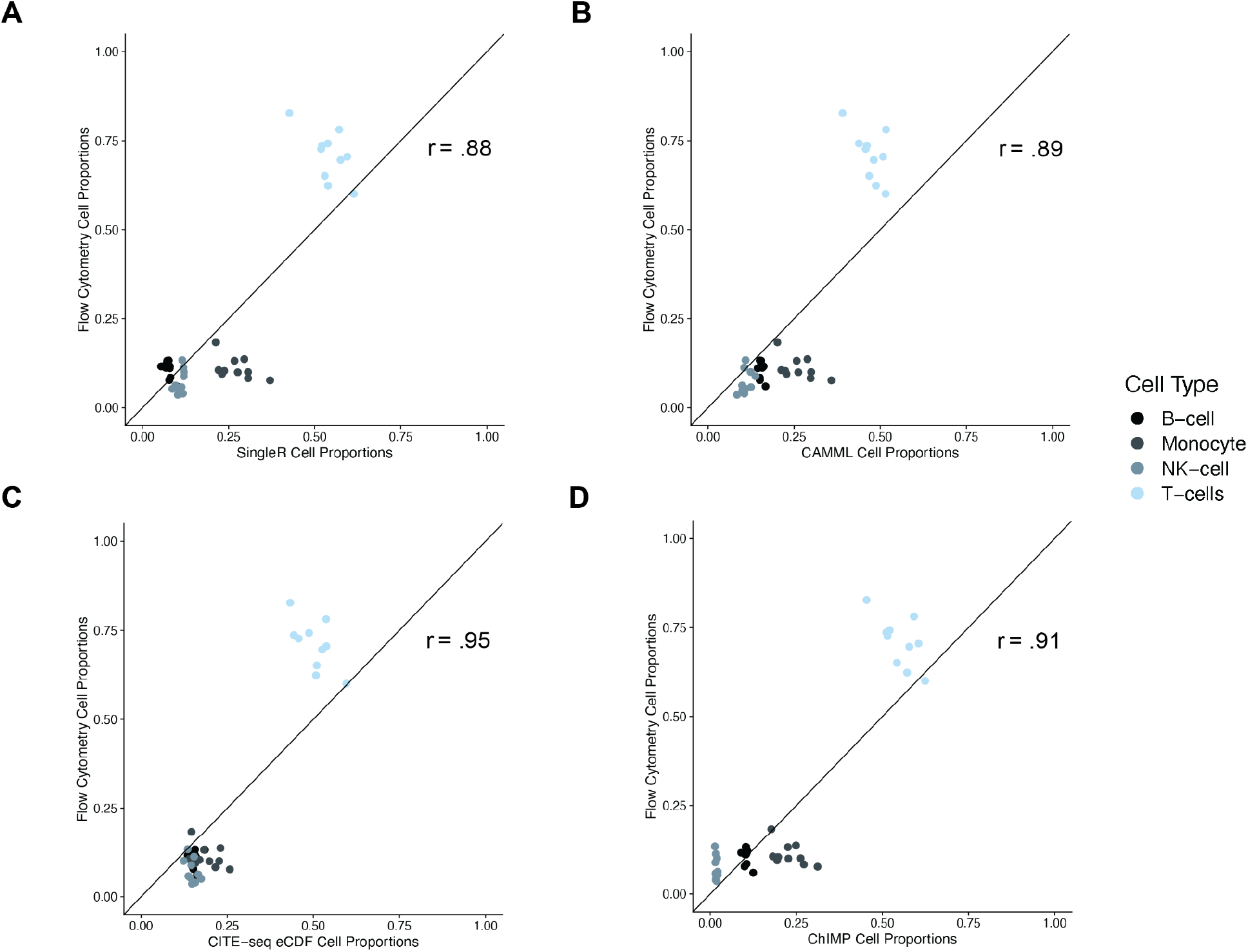
Flow proportions versus SingleR, CAMML, CITE-seq eCDF, and ChIMP. The proportions of cell types identified in the joint scRNA-seq and CITE-seq samples from Lawlor, et al.^28^ by **A**. SingleR, **B**. single-label CAMML, **C**. CITE-seq eCDF, and **D**. ChIMP plotted against the proportions of cell types found in the flow cytometry samples. Each plotted pair of flow-based and scRNA-seq/CITE-seq-based cell proportions were computed using the same tissue sample that was divided prior to FACs and scRNA-seq/CITE-seq analysis.

These results highlight two important considerations:

1. It begs the question of the defining characteristics of a cell type. Given that cell sorting by surface markers has been a mainstay of cell biology research, especially the analysis of immune cells, its use for cell typing of single cell data seems particularly vital. However, in cases where cell phenotype differs from cell surface protein markers, the exclusion of transcriptomic data may give an incomplete picture of cell state and function. For example, a CD8 T cell can be naive, activated, or exhausted, all which have important and distinct implications for a biological system. However, according to cell surface abundance of the CD8 protein, these cells may be indistinguishable. While the CITE-seq eCDF method is the most consistent with previous bulk cell typing methods, the potential information gleaned from combining trancromptomic and surface protein abundance presents a unique and novel direction for cell typing of single cell data.
2. It highlights the challenge of finding a gold standard method for benchmarking cell typing methods that is biased towards neither transcriptomic cell typing nor surface protein cell typing. To evaluate cell typing that is not biased towards a specific omics modality, we utilized a metric, mSDI, previously developed in our lab to characterize cell type confidence independent of a ground truth.

### 2. Hao scRNA-seq/CITE-seq dataata and entropy analysis

Given the aforementioned difficulties with unbiased benchmarking of ChIMP, we decided to evaluate the entropy of cell types by computing the mSDI (Eq. 1) value for each cell in the Hao, et al. joint scRNA-seq/CITE-seq dataset.^31^ Utilizing the mSDI gives a measure of each cell’s cell type entropy, with lower values representing cells that are confidently a single cell type and higher values representing cells that are not confidently identified as any individual cell type. This serves as a useful measure to determine how differentiated a given cell is and how specific a cell typing method is. To evaluate whether cell type entropy was improved with ChIMP relative to other multi-label cell typing approaches, we also computed the mSDI for cell types generated by discretized CITE-seq and CAMML. In this case, we performed cell typing for dendritic cells (DCs), B cells, monocytes, natural killer (NK) cells, and T cells with the following surface markers: CD11c for DCs, CD19 for B cells, CD14 for monocytes, CD56 for NK cells, and either CD4 or CD8 for T cells. Figure 4A-C shows the heatmaps of the scores for discretized CITE-seq, CAMML, and ChIMP respectively. This analysis revealed a marked reduction in the mSDI entropy measure when utilizing ChIMP (median mSDI of 0.047) relative to multi-label CITE-seq (median mSDI of 1.103) and CAMML (median of 0.187) (Figure 4D). In other words, the multi-omics ChIMP method confidently assigned a single cell type to each cell more often than either single-omics component method (i.e, multi-label CITE-seq and CAMML).

**Figure 4.**
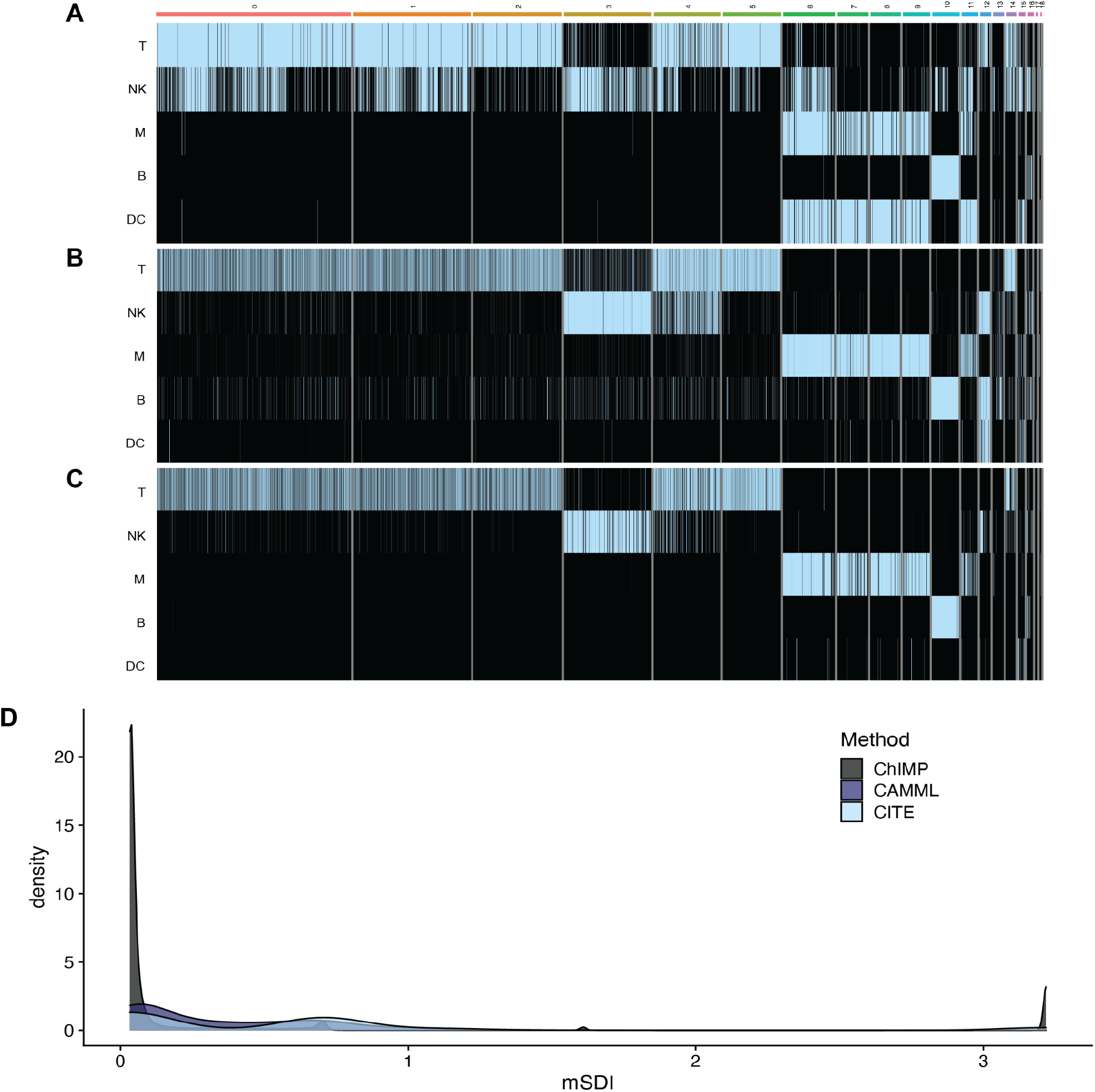
Entropy analysis of ChIMP on PBMCs. **A-C.** Heatmaps of cell type scores (T: T cells, NK: NK cells, M: monocytes, B: B cells, DC: dendritic cells) for discretized CITE-seq, CAMML, and ChIMP respectively. **D**. A density plot of the mSDI scores for each method on the Hao, et al joint scRNA-seq/CITE-seq data.^31^

These results illustrate the potential for ChIMP to serve as a specific and conservative cell type identifier, especially when combined with the promising accuracy results outlined in Figure 3. Depending on the needs of a single cell experiment, having a tool that integrates multiple modalities to make cell type predictions with high specificity could be useful for reducing the risk of type I error and incorrect conclusions. Our entropy analysis of discretized CITE-seq, CAMML, and ChIMP also highlights the notable discordance of cell type calls that can occur based on information from different omics modalities. There may be information lost by utilizing one modality or another, so the nature of these discordant cells and the features that defined their categorization was investigated further.

### 3. MALT scRNA-seq/CITE-seq data

To validate this metric on another dataset and in a different experimental condition, the Li, et al. joint scRNA-seq/CITE-seq dataset of MALT B cell tumors^32^ was scored for discretized CITE-seq, CAMML, and ChIMP, and mSDI was evaluated in each case (Figure 5). We found that, not only did ChIMP again have the lowest median mSDI (0.063 versus 0.613 for CAMML and 0.701 for CITE-seq), but also hat both CAMML and CITE-seq had false positive cell type scores that were ameliorated by the integration of both methods in ChIMP. Specifically, the NK cell marker in the CITE-seq data scores highly in almost every cluster, but there is little indication of a strong NK cell signal with the transcriptomic data, so the signal is removed in ChIMP. Conversely, CAMML scores highly for macrophages in almost every cluster, and while there is some indication of a macrophage population present in clusters 1-3 and 9, the inclusion of CITE-seq removes what is likely to be a false signal in clusters 0 and 4. This false transcriptomic signal is likely due to B cells expressing a gene or pathway that is also associated with macrophages. Indeed, in Figure 7, which shows the top differentially expressed genes, KEGG pathways, and CITE-seq markers in the MALT B cell tumor data, there are genes and pathways that are upregulated in both the B cell and macrophage clusters. This data also illustrates ChIMP’s ability to maintain multiple cell type labels when they are supported by both modalities, as can be visualized with T cells and macrophages in clusters 2-4. It is likely that these clusters either have mixed populations of singleton cells or interacting doublets and ChIMP is still able to identify the potential for multiple cell types in this case–a trait of the original CAMML method that was important to maintain–even with a more conservative method.

**Figure 5.**
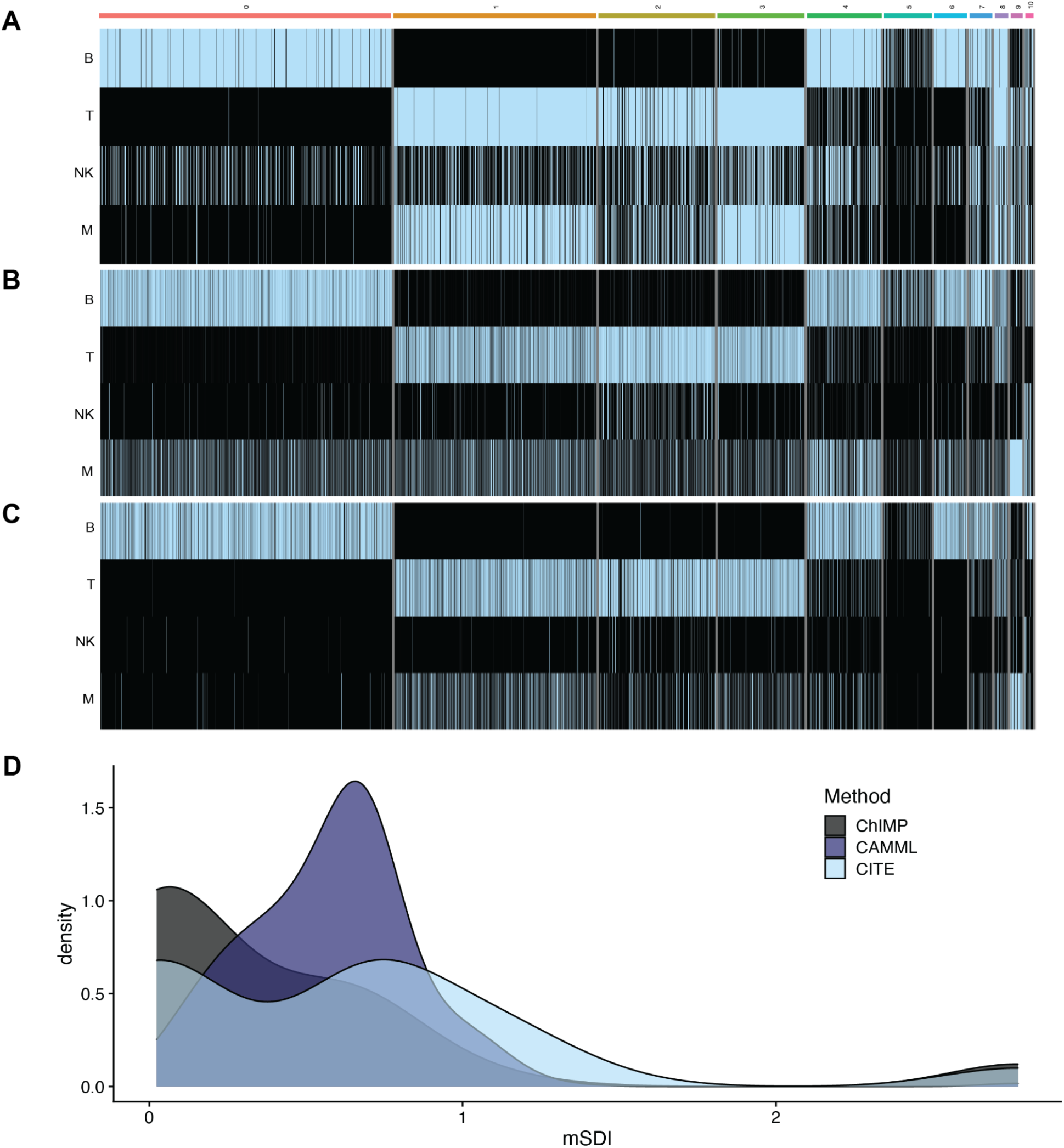
Entropy analysis of ChIMP on MALT B cell tumor data. **A-C.** Heatmaps of cell type scores (B: B cells, T: T cells, M: macrophages, NK: NK cells) for discretized CITE-seq, CAMML, and ChIMP respectively. **D**. A density plot of the mSDI scores for each method on the MALT joint scRNA-seq/CITE-seq data.^32^

### 4. T cell/NK cell discordance in Hao, et al

The most common cell types we found to have discordant classifications in our analysis were T cells and NK cells, especially cells with T cell-like CITE-seq signatures and NK cell-like transcriptomes, as seen in clusters 3 and 4 of the Hao et al. data.^31^ To investigate the phenotypic nature of cells with this juxtaposition, VAM was used to generate cell-level scores for the KEGG^38^ pathways available in MSigDB.^34^ These VAM-based pathway scores, along with the genes and CITE-seq markers in the data, were analyzed for differential expression between clusters using a Wilcoxon Rank Sum test.^39^ The differentially expressed genes, VAM-scored KEGG gene sets, and CITE-seq markers were sorted by logFC for each cluster with the top 3 visualized in a heatmap alongside the ChIMP cell type scores, as shown in Figure 6. We found that the differential expression of pathways involved in cell cytotoxicity, such natural killer cell-mediated cytotoxicity, spliceosome, and proteasome pathways, were highly significant (p <.001) for the discordant cell clusters 3 and 4. This was further supported by the differentially expressed genes, where markers of cytotoxicity, such as perforin and granzyme H, were also highly expressed in both clusters 3 and 4. However, their CITE-seq markers are visibly distinct, with cluster 3 expressing non-T cell markers: CD16 and CD335, and cluster 4 clearly expressing T-cell markers: CD8a and CD8.

**Figure 6.**
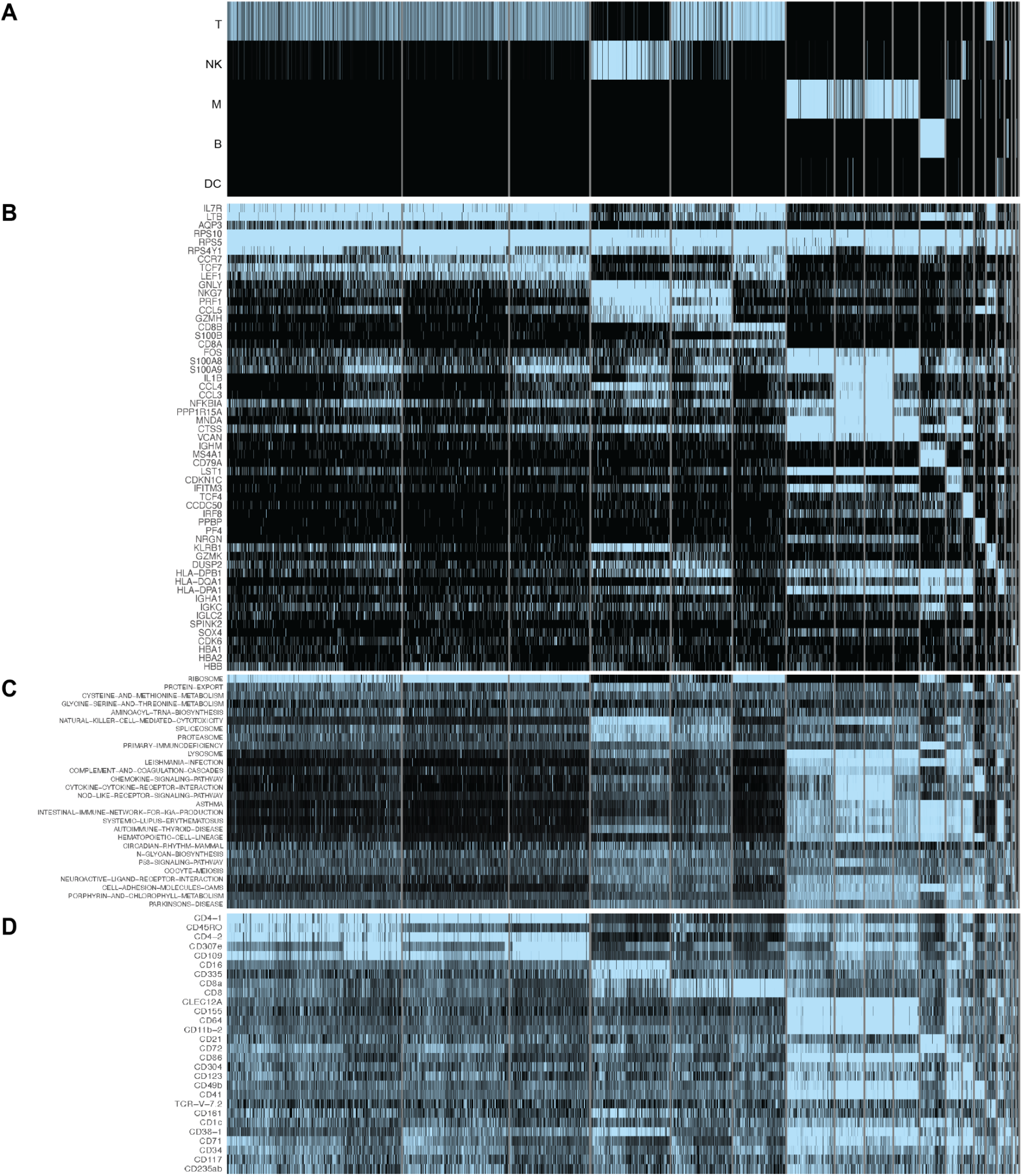
Discordant cell type classifications. Comparison of **A**. the cell type scores for ChIMP, **B**. the top differentially expressed genes across clusters, **C**. the top differentially expressed KEGG pathways across clusters, and **D**. the top differentially expressed CITE-seq markers across clusters in the joint scRNA-seq/CITE-seq data from Hao, et al.^31^

**Figure 7.**
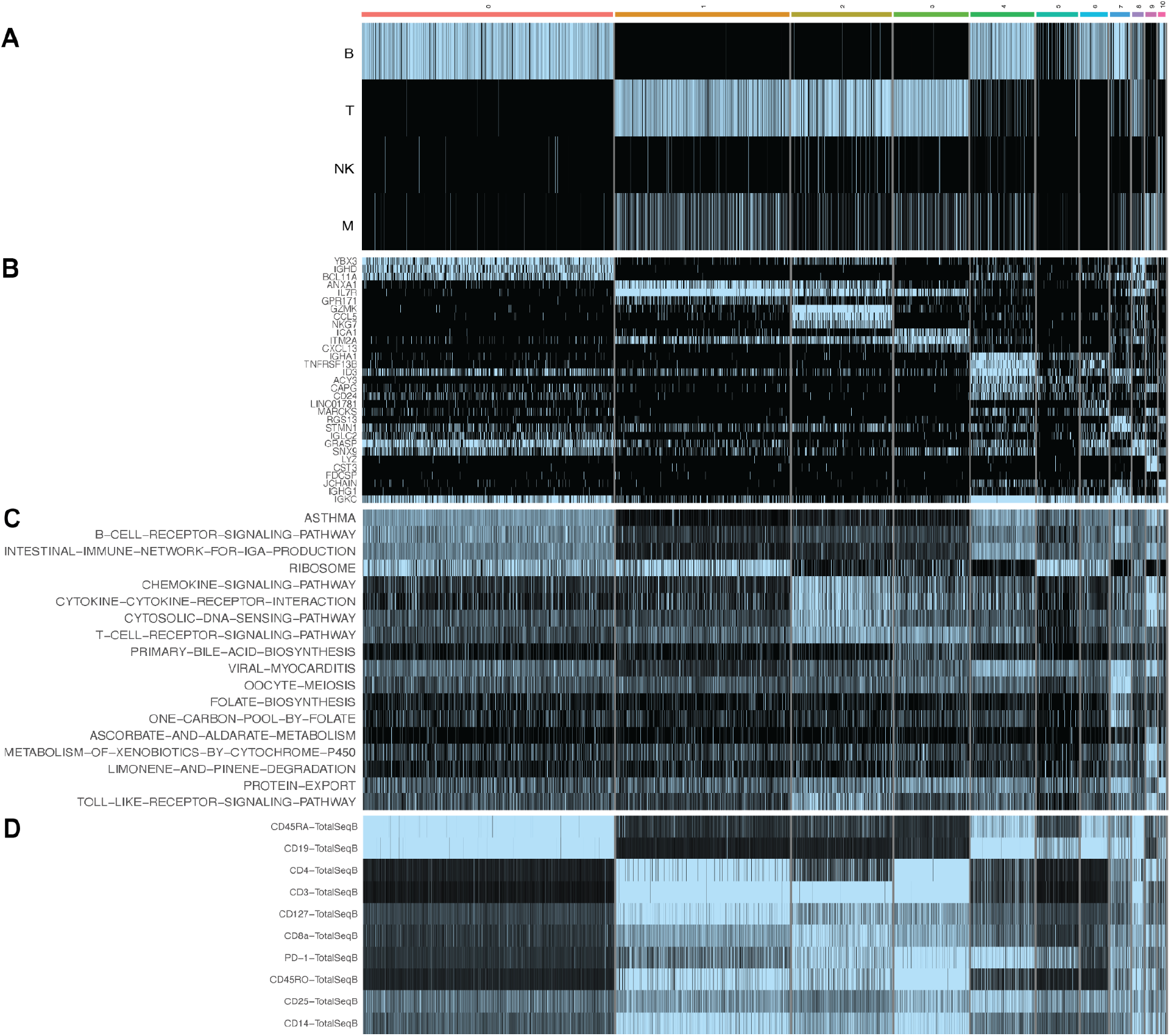
Differential Expression Analysis in Li, et al. MALT B cell tumor data. ^32^ Comparison of **A**. the cell type scores for ChIMP, **B**. the top differentially expressed genes across clusters, **C**. the top differentially expressed KEGG pathways across clusters, and **D**. the top differentially expressed CITE-seq markers across clusters in the joint scRNA-seq/CITE-seq data from Li, et al.^32^

Given that the transcriptomes of cells could not distinguish between T cells and NK cells, the inclusion of surface protein information is important to enable highly specific cell typing. In contrast, the use of surface protein markers alone without gene expression data limits the amount of information that can be gleaned regarding the phenotype of each cell. In this example, clarifying the cell types of discordant cells using CITE-seq gives no context to the highly cytotoxic phenotype occurring in some T cells, which may have a notable influence on the experimental model under investigation. By combining both CITE-seq and scRNA-seq information, ChIMP can consider the important phenotypic and surface protein information in conjunction.

### 5. Customization and inference

ChIMP allows for the integration of scRNA-seq and CITE-seq to be easily customized. The genes used for gene set scoring can be designated and weighted according to an investigator’s needs, either by completely customizing the gene set information or setting the desired differential expression logFC cut-off and gene weights within the CAMML Comprehensive R Archive Network (CRAN) package.^26,37^ Additionally, the markers used for CITE-seq discretization can be customized, both by designating which markers are evaluated for each cell type and by customizing the nature of the required markers. For example, if numerous markers are used for a single cell type, each marker can be required to be in the higher cluster for the CITE-seq score to be 1, or the score can be set to 1 as long as one of the markers is in the higher cluster.

The previously developed CAMML method allows for statistical inference regarding cell type identity via the generated CDF scores.^26,33^ While the ChIMP method alters these scores, multiple hypothesis correction and inference can still be used. Specifically, ChIMP provides more conservative control of the type I error rate than CAMML, i.e., the CAMML-generated p-values are unaltered when the discretized CITE-seq score is 1 or are set to 1 when the CITE-seq score is 0. This correction does not change the inferential capabilities of the scores, but rather makes the evaluation more stringent, i.e., ChIMP reduces the type I error rate at the expense of lower power.

### Conclusion

The advent of single cell genomic profiling has created unique opportunities for tissue characterization, particularly in diseased states.^2–7^ However, analysis of this new single cell data is limited by increased noise and significant sparsity. Investigating the cell types present in these datasets is an ongoing challenge that requires compensating for these data quality challenges.^18,19^ To accomplish this while also addressing the tendency for cell types to occur on a continuum, we developed CAMML, a gene-set based multi-label cell typing tool for scRNA-seq data.^26^ CAMML performs well compared to existing single-label methods and has the added benefit of identifying cells that do not differentiate well into a single category, making it flexible for the analysis of intermediate states, stemness, and multiplets.^26^

With the increasing utilization of multi-omic single cell assays, we extended CAMML to support cell typing of joint scRNA-seq/CITE-seq data data to more confidently and conservatively identify cell types. This new technique, CAMML with the Integration of Marker Proteins (ChIMP), performs multi-label cell typing that accounts for both the expression of cell type transcriptional signatures and abundance of cell type surface protein markers. When comparing ChIMP to ground truth methods of cell typing, ChIMP performs comparably with existing cell-level methods for cell type identification. Furthermore, ChIMP successfully eliminates false positives that occur when each omics modality is used in insolation, resulting in a lower multi-label entropy score while maintaining multiple cell type labels when supported by both modalities. ChIMP is a conservative method for multi-label cell typing of joint scRNA-seq/CITE-seq data based on a statistical model that is robust, transparent and easily customizable.

## Acknowledgements

This work was funded by National Institutes of Health grants R21CA253408, P20GM130454 and P30CA023108. This work was also funded by the Burroughs-Wellcome Fund Big Data in the Life Sciences Training Program. We would like to acknowledge the supportive environment at the Geisel School of Medicine at Dartmouth where this research was performed.

## References

1. Hanahan, D. & Coussens, L. M. Accessories to the crime: functions of cells recruited to the tumor microenvironment. Cancer Cell 21, 309–322 (2012).

2. Azizi, E. et al. Single-Cell Map of Diverse Immune Phenotypes in the Breast Tumor Microenvironment. Cell 174, 1293–1308.e36 (2018).

3. Coffelt, S. B. & de Visser, K. E. Immune-mediated mechanisms influencing the efficacy of anticancer therapies. Trends Immunol. 36, 198–216 (2015).

4. Davidson, S. et al. Single-Cell RNA Sequencing Reveals a Dynamic Stromal Niche That Supports Tumor Growth. Cell Rep. 31, 107628 (2020).

5. Jang, B.-S., Han, W. & Kim, I. A. Tumor mutation burden, immune checkpoint crosstalk and radiosensitivity in single-cell RNA sequencing data of breast cancer. Radiother. Oncol. 142, 202–209 (2020).

6. Qiu, S.-Q. et al. Tumor-associated macrophages in breast cancer: Innocent bystander or important player? Cancer Treat. Rev. 70, 178–189 (2018).

7. Wagner, J. et al. A Single-Cell Atlas of the Tumor and Immune Ecosystem of Human Breast Cancer. Cell 177, 1330–1345.e18 (2019).

8. Wang, L. et al. Single-Cell Map of Diverse Immune Phenotypes in the Metastatic Brain Tumor Microenvironment of Non Small Cell Lung Cancer. bioRxiv (2019) doi:10.1101/2019.12.30.890517.

9. Zhang, Q. et al. Landscape and Dynamics of Single Immune Cells in Hepatocellular Carcinoma. Cell 179, 829–845.e20 (2019).

10. Tuit, S. et al. Transcriptional Signature Derived from Murine Tumor-Associated Macrophages Correlates with Poor Outcome in Breast Cancer Patients. Cell Rep. 29, 1221–1235.e5 (2019).

11. Li, H. et al. Dysfunctional CD8 T Cells Form a Proliferative, Dynamically Regulated Compartment within Human Melanoma. Cell 176, 775–789.e18 (2019).

12. Newman, A. M. et al. Robust enumeration of cell subsets from tissue expression profiles. Nat. Methods 12, 453–457 (2015).

13. Gong, T. & Szustakowski, J. D. DeconRNASeq: a statistical framework for deconvolution of heterogeneous tissue samples based on mRNA-Seq data. Bioinforma. Oxf. Engl. 29, 1083–1085 (2013).

14. Li, X. & Wang, C.-Y. From bulk, single-cell to spatial RNA sequencing. Int. J. Oral Sci. 13, 1–6 (2021).

15. Chen, G., Ning, B. & Shi, T. Single-Cell RNA-Seq Technologies and Related Computational Data Analysis. Front. Genet. 10, 317 (2019).

16. Tang, F. et al. mRNA-Seq whole-transcriptome analysis of a single cell. Nat. Methods 6, 377–382 (2009).

17. Hay, S. B., Ferchen, K., Chetal, K., Grimes, H. L. & Salomonis, N. The Human Cell Atlas bone marrow single-cell interactive web portal. Exp. Hematol. 68, 51–61 (2018).

18. Kolodziejczyk, A. A., Kim, J. K., Svensson, V., Marioni, J. C. & Teichmann, S. A. The Technology and Biology of Single-Cell RNA Sequencing. Mol. Cell 58, 610–620 (2015).

19. Haque, A., Engel, J., Teichmann, S. A. & Lönnberg, T. A practical guide to single-cell RNA-sequencing for biomedical research and clinical applications. Genome Med. 9, 75 (2017).

20. Diaz-Mejia, J. J. et al. Evaluation of methods to assign cell type labels to cell clusters from single-cell RNA-sequencing data. F1000Research 8, 296 (2019).

21. Kiselev, V. Y., Andrews, T. S. & Hemberg, M. Challenges in unsupervised clustering of single-cell RNA-seq data. Nat. Rev. Genet. 20, 273–282 (2019).

22. Tirosh, I. et al. Dissecting the multicellular ecosystem of metastatic melanoma by single-cell RNA-seq. Science 352, 189–196 (2016).

23. Satija, R., Farrell, J. A., Gennert, D., Schier, A. F. & Regev, A. Spatial reconstruction of single-cell gene expression data. Nat. Biotechnol. 33, 495–502 (2015).

24. de Kanter, J. K., Lijnzaad, P., Candelli, T., Margaritis, T. & Holstege, F. C. P. CHETAH: a selective, hierarchical cell type identification method for single-cell RNA sequencing. Nucleic Acids Res. 47, e95–e95 (2019).

25. Aran, D. et al. Reference-based analysis of lung single-cell sequencing reveals a transitional profibrotic macrophage. Nat. Immunol. 20, 163–172 (2019).

26. Schiebout, C. & Frost, H. R. CAMML: Multi-Label Immune Cell-Typing and Stemness Analysis for Single-Cell RNA-sequencing. Pac. Symp. Biocomput. (2022).

27. Stoeckius, M. et al. Simultaneous epitope and transcriptome measurement in single cells. Nat. Methods 14, 865–868 (2017).

28. Lawlor, N. et al. Single Cell Analysis of Blood Mononuclear Cells Stimulated Through Either LPS or Anti-CD3 and Anti-CD28. Front. Immunol. 12, 636720 (2021).

29. McInnes, L., Healy, J. & Melville, J. UMAP: Uniform Manifold Approximation and Projection for Dimension Reduction. ArXiv180203426 Cs Stat (2020).

30. Blondel, V. D., Guillaume, J.-L., Lambiotte, R. & Lefebvre, E. Fast unfolding of communities in large networks. J. Stat. Mech. Theory Exp. 2008, P10008 (2008).

31. Hao, Y. et al. Integrated analysis of multimodal single-cell data. Cell 184, 3573–3587.e29 (2021).

32. Li, L. et al. Improved integration of single-cell transcriptome and surface protein expression by LinQ-View. Cell Rep. Methods 1, 100056 (2021).

33. Frost, H. R. Variance-adjusted Mahalanobis (VAM): a fast and accurate method for cell-specific gene set scoring. Nucleic Acids Res. 48, e94–e94 (2020).

34. Liberzon, A. et al. Molecular signatures database (MSigDB) 3.0. Bioinformatics 27, 1739–1740 (2011).

35. Robinson, M. D., McCarthy, D. J. & Smyth, G. K. edgeR: a Bioconductor package for differential expression analysis of digital gene expression data. Bioinformatics 26, 139–140 (2010).

36. Mabbott, N. A., Baillie, J. K., Brown, H., Freeman, T. C. & Hume, D. A. An expression atlas of human primary cells: inference of gene function from coexpression networks. BMC Genomics 14, 632 (2013).

37. R Core Team. R: A Language and Environment for Statistical Computing. (R Foundation for Statistical Computing).

38. Kanehisa, M. & Goto, S. KEGG: Kyoto Encyclopedia of Genes and Genomes. Nucleic Acids Res. 28, 27–30 (2000).

39. Wilcoxon, F. Individual Comparisons by Ranking Methods. Biom. Bull. 1, 80–83 (1945).

